# Molecular basis of Gabija anti-phage supramolecular assemblies

**DOI:** 10.1101/2023.08.07.552356

**Authors:** Xiao-Yuan Yang, Zhangfei Shen, Qingpeng Lin, Tian-Min Fu

**Affiliations:** Department of Biological Chemistry and Pharmacology, The Ohio State University, Columbus, OH 43210, USA; The Ohio State University Comprehensive Cancer Center, Columbus, OH 43210, USA; Program of OSBP, The Ohio State University, Columbus, OH 43210, USA

## Abstract

As one of the most prevalent anti-phage defense systems in prokaryotes, Gabija consists of a Gabija protein A (GajA) and a Gabija protein B (GajB). The assembly and function of the Gabija system remain unclear. Here we present cryo-EM structures of GajA and the GajAB complex, revealing tetrameric and octameric assemblies, respectively. In the center of the complex, GajA assembles into a symmetric tetramer, which recruits two sets of GajB dimer at opposite sides of the complex, resulting in a 4:4 GajAB supramolecular complex for anti-phage defense. Further biochemical analysis showed that GajA alone is sufficient to cut double-stranded DNA and plasmid DNA, which can be inhibited by ATP. Unexpectedly, the GajAB displays enhanced activity for plasmid DNA, suggesting a role of substrate selection by GajB. Together, our study defines a framework for understanding anti-phage immune defense by the GajAB complex.

## Introduction

To mitigate phage infections, bacteria have evolved highly diverse anti-phage immune systems ^1^. Though some bacterial immune systems like CRISPR-Cas and CBASS have been extensively studied ^2, 3^, many newly identified systems remain unexplored ^4–6^. The study of bacteria immune system not only offers evolutionary perspectives on immune systems but also provides invaluable tools for biomedical research and disease treatment.

As a newly identified bacteria immune system, bacterial Gabijia defense system exists in at least 8.5% of sequenced genomes with two components, GajA and GajB^4, 7^. GajA was shown to be an endonuclease that can recognize specific DNA sequence^8^. GajB was predicted to be a UvrD-like helicase^9^. However, whether and how GajA and GajB assemble into a complex for anti-phage defense remains unclear.

Here, we present the cryo-EM structures of GajA alone and the GajAB complex. GajA assembles into a tetramer via interactions mediated by both the ATPase domain and the nuclease domain. We also revealed that the GajA and GajB assemble into a heteromeric octamer with four molecules of GajA and four molecules of GajB, which is critical for the anti-phage defense. Given many other supramolecular assemblies identified in bacterial immunity, we propose that supramolecular assemblies may represent a unified mechanism in bacterial immune defense.

## Results

### Structure of GajA

To biochemically characterize GajA, we expressed and purified GajA in *E. coli* BL21(DE3) (Extended data Fig. 1a-b). The elution volume of GajA on gel filtration indicated that GajA formed an oligomer (Extended data Fig. 1a). To reveal the assembly of GajA, we employed cryo-EM single particle analysis to determine the structure of GajA. However, the GajA had severe orientation preference problem on grids, leading to a reconstruction with an apparent resolution of 2.9 Å but poor densities (Extended data Fig. 1c-g). To resolve this issue, we optimized conditions for grids preparation and eventually obtained a cryo-EM structure of GajA with a resolution of 3.2 Å by collecting a dataset using a grid with relatively thicker ice (Extended data Fig. 1a and 2a-e, table 1). Although the apparent resolution is lower from reconstruction of images with thicker ice, the EM densities have been significantly improved in comparison to the 2.9 Å structure (Extended data Fig. 1g and 2f). The cryo-EM structure of GajA revealed a symmetric tetrameric assembly with dimensions of 175 Å × 115 Å × 50 Å (Fig. 1a-b).

**Fig. 1.**
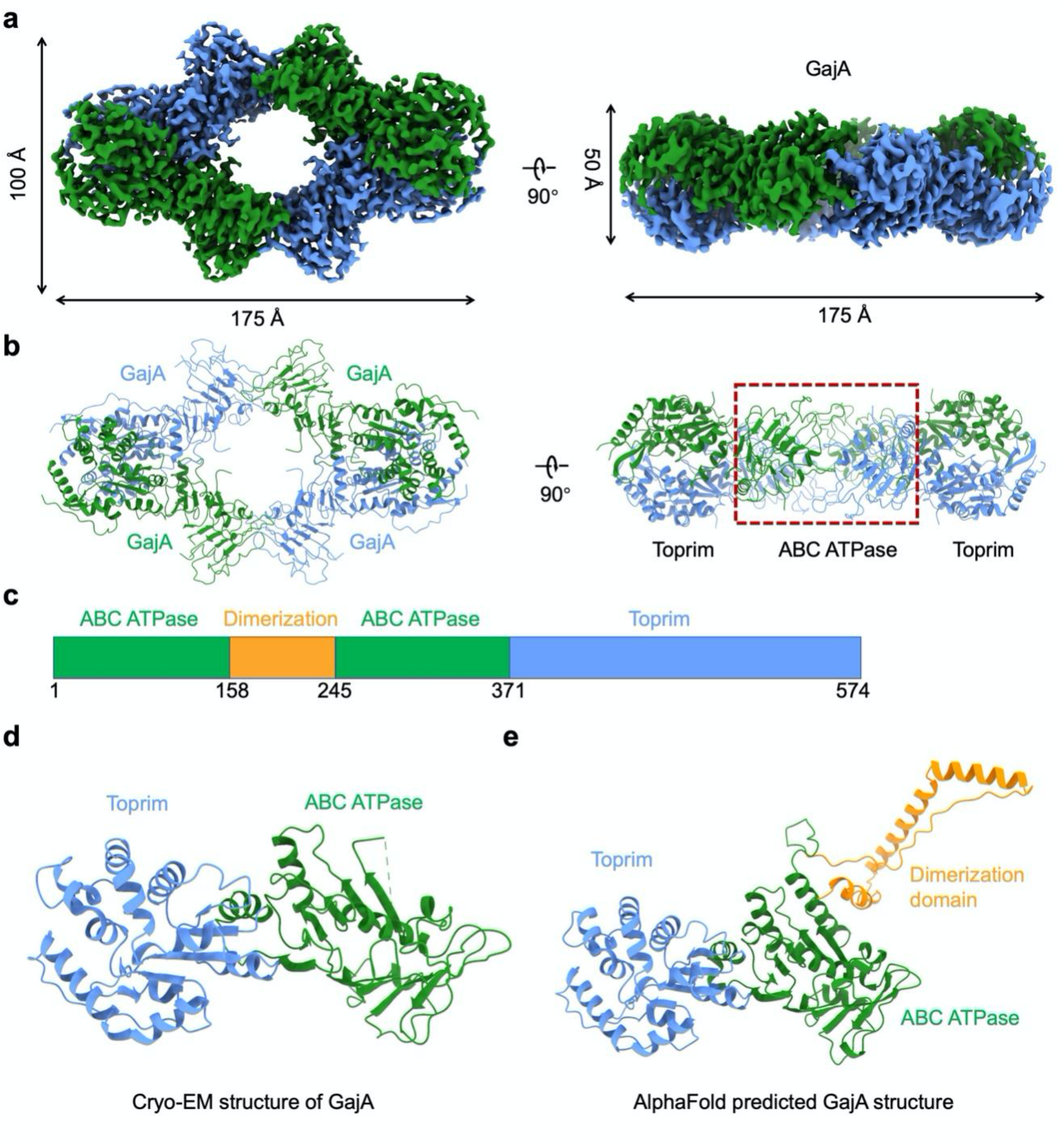
Cryo-EM structure of GajA. **a, b,** Cryo-EM density map (**a**) and ribbon diagrams (**b**) of GajA tetramer. **c,** Domain architecture of GajA. The ABC ATPase domain is indicated in green, dimerization domain in orange, and Toprim in blue. **d, e,** Ribbon diagram of a GajA protomer determined by cryo-EM reconstruction (**d**) or AlphaFold prediction (**e**) with domains colored as in **c**.

**Fig. 2.**
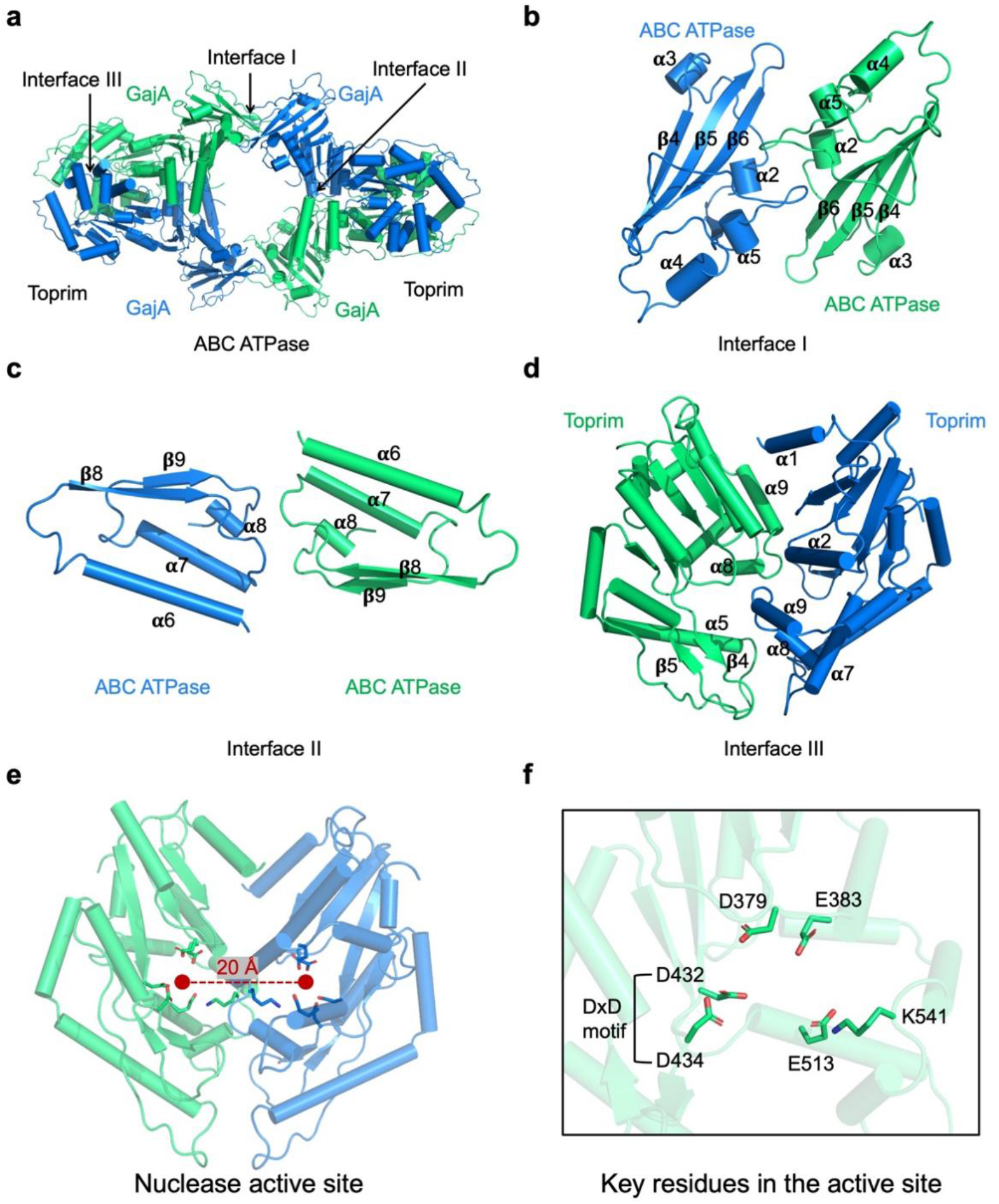
Assembly of GajA. **a,** Assembly of tetrameric GajA with three key interfaces indicated, which are denoted as Interface I, II, and III. **b,** Details of interface I mediated by the first haves of ABC ATPase domains with secondary structures indicated. **c,** Details of interface II mediated by the second halves of ABC ATPase domains with secondary structure indicated. **d,** Details of interface III mediated by the toprim domains with secondary structure indicated. **e,** Catalytic center of the toprim domains. Distance between the active sites of dimeric toprim domains are highlighted. **f,** Key residues in the catalytic center of the toprim domain that are highlighted in sticks.

Each protomer of GajA is composed of an N-terminal ATPase domain that is divided into two halves by an inserted dimerization domain, and a C-terminal nuclease domain (Fig. 1c-e). The N-terminal ATPase domain is composed of a 11-stranded mixed-paralleled β-sheet, sandwiching α1 and surrounded by α2-α8 (Fig. 1d and Extended data Fig. 3a). Structural comparison revealed that the GajA ATPase domain resembles the canonical <u>A</u>TP-<u>b</u>inding <u>c</u>assette (ABC) ATPase proteins with a highly conserved ATP binding site ^10, 11^ (Extended data Fig. 3b). The GajA C-terminal domain folds as a Toprim (topoisomerase-primase) domain with a central four-stranded parallel β-sheet surrounded by α-helices ^12, 13^ (Fig. 1d and Extended data Fig. 3c). Both the N-terminal ATPase domain and the C-terminal Toprim domain were clearly resolved in our cryo-EM structure (Fig. 1d). In contrast, the dimerization domain, predicted to consist of three α-helices by AlphaFold ^14^, was invisible, indicating the flexibility of this domain (Fig. 1d-e).

### Assembly of GajA tetramer

The tetramerized GajA is arranged as a dimer of dimer with three types of interfaces, denoted as Type I, Type II, and Type III interfaces (Fig. 2a). The Type I interface, located at the center of GajA tetramer, is mediated by the first half of the ATPases domain with a buried surface area of 609 Å^2^ (Fig. 2b). Detailed analysis revealed that hydrophobic interactions dominate the formation of type I interface (Extended data Fig. 4a). In contrast, both hydrophobic and hydrophilic residues in the second half of the ATPase domain contribute to the formation of Type II interface that is characterized by a buried surface area of 325 Å^2^ (Fig. 2c and Extended data Fig. 4b). Additionally, Toprim domains are arranged along a 2-fold axis to form the Type III interface with an extensively buried surface area of 1,471 Å^2^ (Fig. 2d and Extended data Fig. 4c). Together, interactions of these three interfaces govern the assembly of a 4-fold symmetric GajA.

### Active site of GajA

The Toprim domains in GajA tetramer, which belong to a class of DNA endonucleases known as OLD (overcoming lysogenization defect) ^12^, dimerize at opposite sides of the elongated tetramer. Analysis of the two proximal Toprim domains reveals that the two active sites are about 20 Å away from each other, suggesting that the two active sites work independently (Fig. 2e). Similar to other OLD nucleases^12, 13^, the active site of GajA is composed of a conserved DxD motif between α3 and β3, an invariant glutamate following β2, and an invariant glycine in the α1-β1 loop (Fig. 2f). Studies on BpOLD suggested a two-metal catalysis mechanism^13^, which may be shared by GajA due to the structural similarity of the active sites between GajA and BpOLD (Extended data Fig. 4d). The conserved DxD motif and D379 may coordinate a magnesium while the other magnesium may be coordinated by E513 and E383^13, 15^ (Extended data Fig. 4d). Consistently, D379A mutation abolished the nuclease activity of GajA, highlighting the significant role of D379 in catalysis^8^.

### Structure of GajAB complex

To understand the assembly of GajAB, we further reconstituted the GajAB complex and determined a 3.0 Å cryo-EM structure of the complex (Fig. 3a-b and Extended data Fig. 5a-c, table 1). The cryo-EM structure of GajAB reveals a 4:4 assembly of GajA and GajB with a dimension of 175 Å×145 Å×95 Å, contrasting with a previous assumption that GajA and GajB form a complex with variable stoichiometries^9^ (Fig. 3a-b). In the GajAB complex, the tetrameric GajA is decorated by a pair of GajB dimers at either end (Fig. 3a-b).

**Fig. 3.**
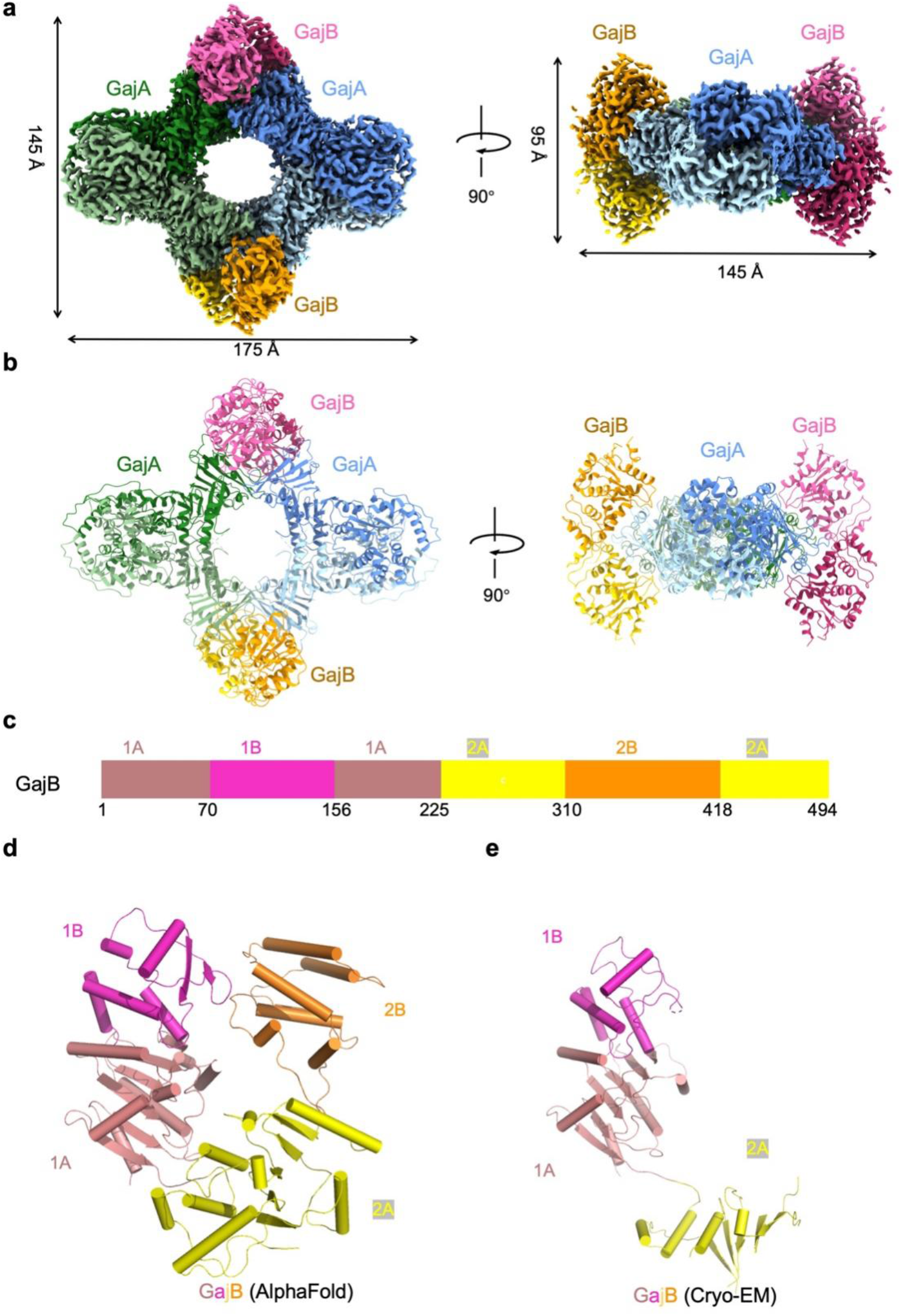
Structure of the GajAB complex. **a, b,** Cryo-EM density map (**a**) and ribbon diagrams (**b**) of the GajAB complex with GajA in cold colors and GajB in warm colors. **c,** Domain architecture of GajB. The 1A domain is indicated in pink, 1B domain in magenta, 2A in yellow, and 2B in orange. **d, e,** Ribbon diagram of a GajB protomer predicted by AlphaFold (**d**) or determined by cryo-EM reconstruction (**e**) with domains colored as in **c**.

GajB is composed of four structural domains 1A, 1B, 2A, and 2B, resembling superfamily 1A helicase proteins like UvrD, PcrA, and Rep that function to unwind and translocate DNA ^16–18^ (Fig. 3c-d and Extended data Fig. 6a). In our cryo-EM structure, domains 1A, 1B, and part of the 2A are visualized, while domain 2B is completely absent (Fig. 3e). Sequence alignment revealed that GajB contains the eight sequence motifs in domains 1A and 2A of UvrD, which have been identified to be involved in ATP binding^18^ (Extended data Fig. 6b). These features indicated that GajB is capable of hydrolyzing ATP. Consistently, ATPase activities are detected in GajB in the presence of DNA^9^, indicating DNA substrates are required to stimulate the ATPase activity of GajB. Structural comparison to UvrD revealed that domain 2A in GajB is not well positioned to coordinate ATP (Extended data Fig. 6c). As such, conformational changes are required for GajB to bind and hydrolyze ATP upon DNA binding. The structure of UvrD in complex with ds-ss DNA junction revealed that the single-strand DNA binds to domains 1A and 2A at their interfaces with 1B and 2B while the DNA duplex is coordinated by domains 1B and 2B^18^. Comparisons with UvrD demonstrate that GajB contains all the key residues for coordinating ssDNA, suggesting that GajB may use a similar strategy for binding ssDNA (Extended data Fig. 6d). In contrast, the domain 2B in GajB is much smaller than that of UvrD and lacks key residues for coordinating dsDNA, raising questions whether GajB can efficiently bind to dsDNA (Extended data Fig. 6a and 6e).

### Mechanism of GajAB assembly

The structure of the GajAB complex revealed that the GajA recruits a pair of GajB molecules via its ATPase domain (Fig. 4a-c). Each GajB molecule interacts with two GajA molecules (*cis* and *trans*) that form a head-to-head dimer via their ATPase domain (Fig. 4b-c). The interaction between GajB and GajA in *cis* is quite extensive with a buried surface area of 755 Å^2^, which is dominated by interactions between GajB 1B domain and GajA ATPase domain (Fig. 4b and Extended Data Fig. 7a-b). Both hydrophobic and hydrophilic residues form an extensive network to dock GajB 1B domain onto the ATPase domain of GajA, positioning the GajB 1A domain adjacent to the GajA ATPase domain with relatively weaker interactions (Extended Data Fig. 7a-b). In contrast, the interactions between GajB and GajA in *trans* are much weaker with a buried surface area of 140 Å^2^, which is mediated by GajB 1A domain and GajA ATPase domain (Fig. 4c and Extended Data Fig. 7c). Additionally, the two paired GajB molecules form relatively weak interactions with each other with a buried surface area of 380 Å^2^, which is dominated by hydrophilic residues in the 1B domain of GajB (Fig. 4d and Extended Data Fig. 7d). Collectively, these three interfaces together with the extensive interactions among tetrameric GajA form the basis of GajAB supramolecular complex assembly.

**Fig. 4.**
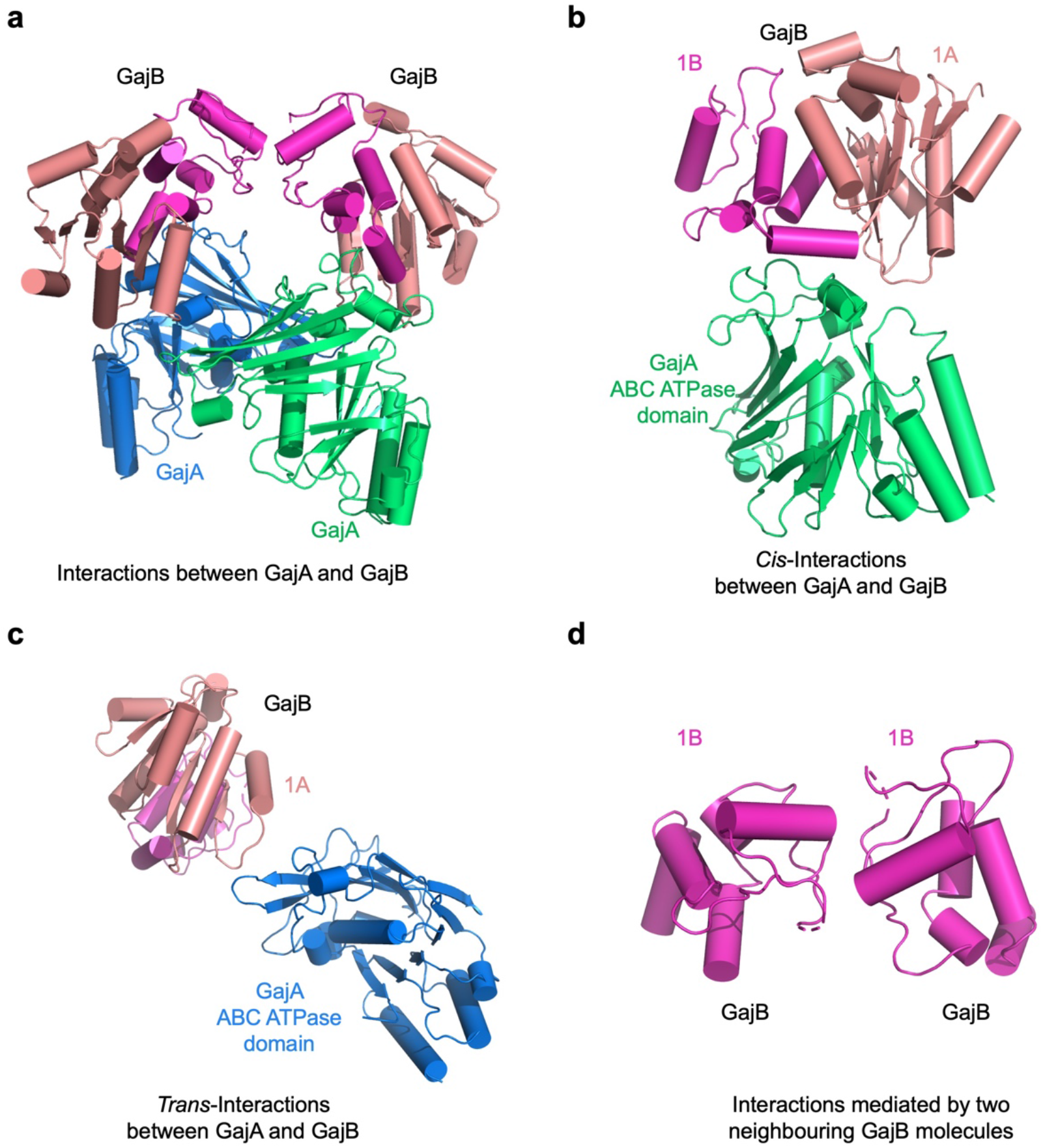
Assembly of GajAB. **a,** Assembly of GajAB with a dimeric GajA (green and blue) engaged with two GajB protomers (pink and magenta). **b,** *Cis*-interactions mediated by GajA ATPase domain and GajB. **c,** *Trans*-interactions mediated by GajA ATPase domain and GajB. **d,** Interactions between two neighboring GajB protomers, which are mediated by the 1B domain of GajB.

### Nuclease activities and anti-phage defense of GajAB complex

Consistent with previous studies^8, 9^, GajA has nuclease activities in the presence of Mg^2+^ and is capable of cleaving dsDNA while GajB has no nuclease activities (Fig. 5a-b). Unexpectedly, the complex of GajAB displayed similar nuclease activities towards dsDNA as GajA (Fig. 5c). Contrasting a previous study^8^, we found that both GajA and GajAB are capable of cleaving plasmid pUC19 (Fig. 5d-e). Moreover, GajAB displayed higher nuclease activities towards pUC19 than GajA alone, highlighting the importance of GajAB complex assembly in effectively cleaving plasmid DNA (Fig. 5d-e). Further phage resistance assay showed that the GajAB complex is more effective in anti-phage defense compared to GajA or GjaB alone (Fig. 5f). In addition, we also showed that the nuclease activity of GajAB is essential for anti-phage defense (Fig. 5f). As such, the supramolecular complex assembly of GajAB is critical for anti-phage defense.

**Fig. 5.**
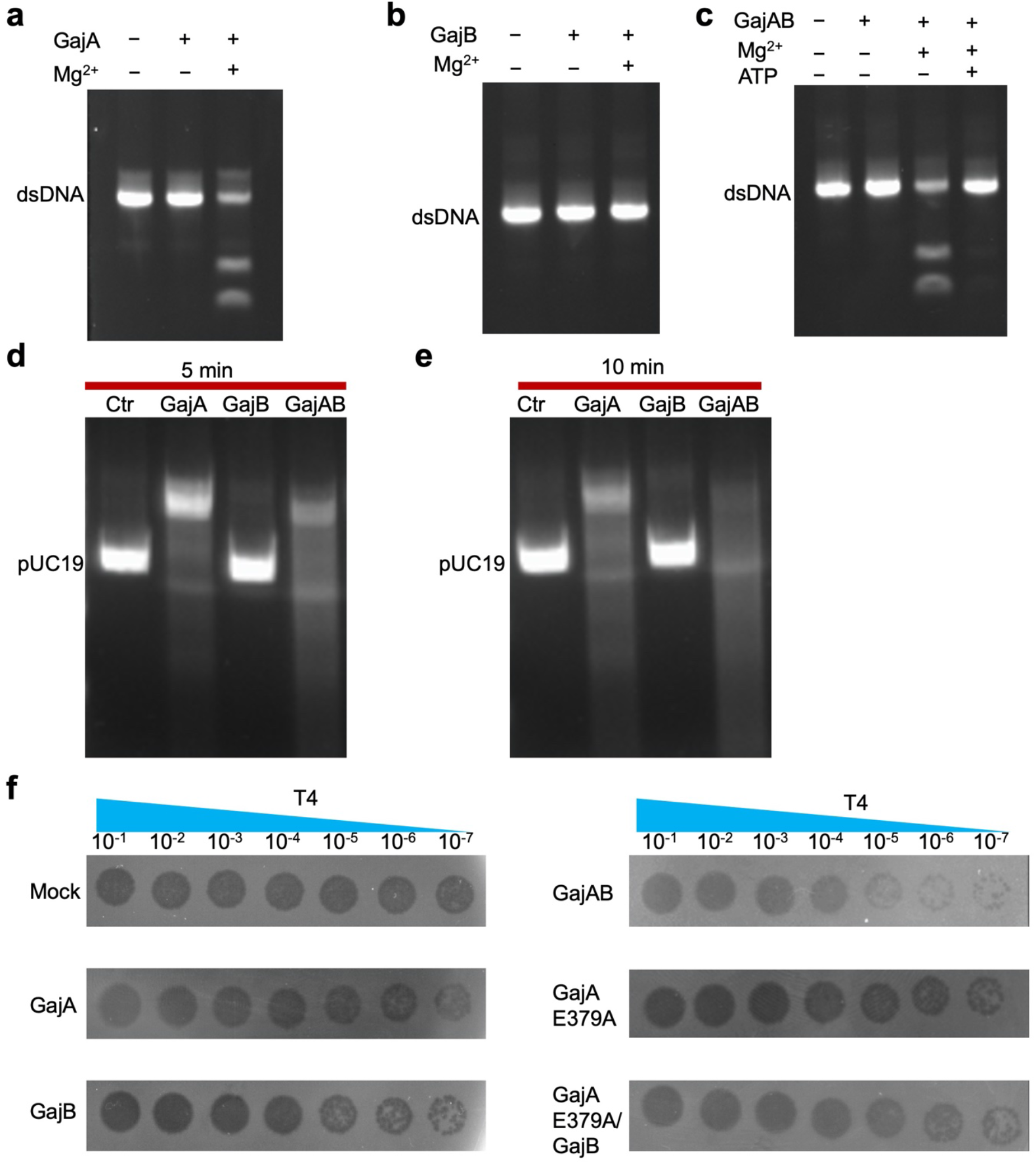
Anti-phage defense of GajAB. **a,** dsDNA cleavage by GajA in the presence of magnesium. **b,** dsDNA cannot be processed by GajB. **c,** GajAB cleaves dsDNA in the presence of magnesium, which can be inhibited by ATP. **d, e,** pUC19 plasmids were processed by GajA, GajB, and GajAB for 5 minute (**d**) and 10 minutes (**e**) at room temperature, respectively. GajAB displayed higher activities than GajA, underscoring the importance of GajB in promoting the catalytic activity of GajA. **f,** Anti-phage defense of GajA, GajB, GajAB, GajA E379A mutant, and the complex of GajA E379A and GajB.

## Discussion

We find that GajA and GajB assemble into a supramolecular complex for anti-phage defense (Fig. 6). Our structural analysis revealed that GajA alone forms a tetramer and further assembles into a heteromeric octamer with GajB (Fig. 6). Moreover, GajA alone has nuclease activity while GajB adopts similar fold as UvrD that can bind DNA substrates. No obvious conformational changes have been observed in GajA upon binding to GajB. As such, we propose that GajB may function to assist GajA to better recognize its substrates in vivo rather than directly promote the catalytic activities of GajA. Consistent with this assumption, we found that the complex of GajAB displays higher nuclease activities towards plasmids than GajA alone (Fig. 5d and 5e). Additionally, the nuclease activity of GajA is inhibited by ATP. It is possible that DNA substrates or other factors in phages may activate the Gabjia system by relieving the inhibitory effect of ATP.

**Fig. 6.**
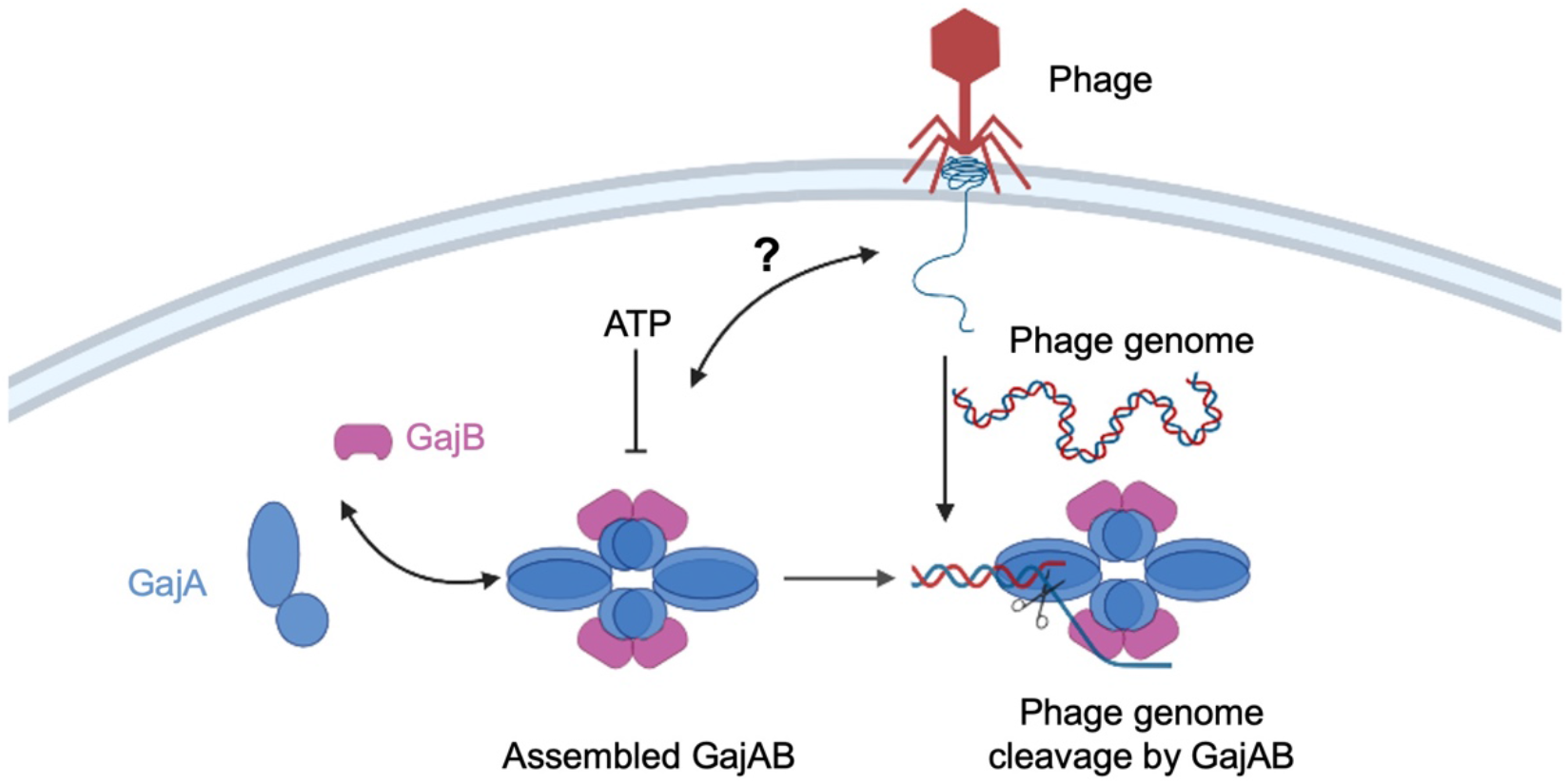
Mechanisms of GajAB assembly and function. A schematic diagram to illustrate mechanisms of GajAB assembly and function.

Supramolecular assembly appears to be an emerging theme in anti-phage immune defense. More and more studies have revealed that bacterial immune systems tend to form large complexes for anti-phage defense ^19–21^. For example, RdrA and RdrB in the RADAR system assemble into a giant assembly with a molecular weight of up to 10 MDa ^19, 20^. Here, we present another example to show the supramolecular assembly by the Gabija system. As both RADAR system and Gabija system contain and oligomerize via ATPase domains, we believe that other ATPase-containing bacterial immune systems may also assemble into supramolecular complexes for anti-phage defense.

## Materials and methods

### Molecular cloning, protein expression and purification

*Bacillus cereus (B. cereus)* GajA (UniProt: J8H9C1) with an N-terminal His×6-tag was cloned into the pET28a vector. *B. cereus* GajB (UniProt: J8HQ06) was inserted into the pETDuet-1 vector with an N-terminal His×6 tag. All the mutants were made through site-direction mutagenesis.

Recombinant plasmids for protein expression were transformed into *E. coli* BL21 (DE3) cells (ThermoFisher Scientific), which were cultured in LB medium containing 50 μg/ml kanamycin at 37°C. When an OD600 of 0.6 - 0.8 was reached, protein expression was induced at 18°C by 0.3 mM IPTG. Cells were harvested after overnight induction (∼16h) and resuspended in lysis buffer (50 mM Tris-HCl pH 8.0, 500 mM NaCl, 10 mM imidazole). After sonication, the supernatant of lysate was collected through centrifugation at 30,000 x g, 4 °C for 50 min. The clarified lysate was loaded onto a pre-equilibrated Ni^2+^-NTA agarose column, and then the column was washed with 30 column volumes (CV) of Ni^2+^-NTA wash buffer (50 mM Tris-HCl pH 8.0, 500 mM NaCl, 10 mM imidazole). The protein was eluted in elution buffer (50 mM Tris-HCl pH 8.0, 150 mM NaCl, 250 mM imidazole, 0.4 mM TCEP). Protein was further purified by size exclusion chromatography using gel-filtration column (Superose 6 increase 10/300 GL, Cytiva, Sigma-Aldrich) in a buffer containing 50 mM Tris-HCl pH 8.0, 150 mM NaCl, and 0.4 mM TCEP.

For the assembly of the GajAB complex, we incubated GajA and GajB with molar ratio of 1:1 on ice for 1 hour followed by further purification via gel filtration.

### Cryo-EM data collection

3 μL sample at 1.8 mg/ml was applied to a glow-discharged Quantifoil R1.2/1.3 400 mesh gold grid (Electron Microscopy Sciences), blotted for 4 s in 100% humidity at 4 °C and plunged into liquid ethane using an FEI Vitrobot Mark IV (Thermo Fisher). All grids were screened using a ThermoFisher Glacios microscope (OSU Center for Electron Microscopy and Analysis).

For GajA tetramer (4A) in thicker ice, 1,364 micrographs were collected using a 300 kV Titan Krios microscope equipped with a K3 direct electron detector (Thermo Fisher) in counting mode with a nominal magnification of 81,000×, and a physical pixel size of 0.899 Å with defocus values ranging from -1.0 to -2.0 μm. For GajA in thin ice, 6,370 images were collected using similar parameters.

For GajAB hetero-complex (4A:4B), 7173 micrographs were collected using a K3 detector with physical pixel size of 1.12 Å. Each micrograph stack contains 40 frames with a total electron dose of 50 e^-^/Å^2^ s.

### Cryo-EM data processing

The detailed flowcharts for data processing of all the datasets were illustrated, respectively (Extended data Fig. 1e, 2c, and 5c). The datasets were imported into cryoSPARC (v4.1.1) implementation of patch motion correction, and patch contrast transfer function (CTF) estimation ^22^. Initial particle picking was done by blob picking to generate initial 2D classes. Representative 2D classes were then selected as templates to pick all the particles for reconstruction.

For GajA tetramer, 849,640 particles were picked and extracted. After two rounds of 2D classification, 583,860 particles were selected and merged for further 3D reconstruction and heterogeneous refinement. The best class of 96,633 particles were selected for further non-uniform refinement with D2 symmetry, resulted in a 3.23 Å map.

For GajAB complex, 6,928,153 particles were picked and extracted. After two rounds of 2D classification, 2,682,203 particles were selected for further ab-initio reconstruction to generate three initial models for further refinement. The best class was selected for further 3D classification and heterogeneous refinement. The final best class of 942,091 particles were selected for non-uniform refinement with C1 symmetry and D2 symmetry, resulted in a 2.98-Å map and a 2.79-Å map, respectively.

All reported resolutions were estimated based on the gold-standard Fourier shell correlation (FSC) = 0.143 criterion ^23^.

### Model building and refinement

Two initial models of Gabija protein A and Gabija protein B were predicted by AlphaFold ^14^, and fitted into the cryo-EM maps of Gabija A tetramer or Gabija AB complex using Chimera^24^. Manual adjustments were done using Coot to yield the final atomic model^25^. Real-space refinement was performed to refine the model against cryo-EM density map with secondary structure and geometry restraints in PHENIX^26^. The all-atom contacts and geometry for the final models were validated by Molprobity^27^. All the structural figures were generated using PyMOL^28^, Chimera^24^, and ChimeraX^29^.

### Nuclease assays

For plasmid DNA, 400 nM protein was incubated with 800 nM substrates in reaction buffer (50 mM Tris-HCl pH 8.0, 150 mM NaCl, and 5 mM MgCl_2_) at 37 °C for 15 min. The final products were separated in 1 % (w/v) Agar LB gel at 100 V in 1 x TAE buffer for 30 mins.

For dsDNA substrates, 400 nM protein were incubated with 200 nM 5’ FAM-labeled nucleic acids substrate in reaction buffer (50 mM Tris-HCl pH 8.0, 150 mM NaCl, and 5 mM MgCl_2_) at 37 °C for 10 min. Products were separated using 12% PAGE gel at 150 V in 1 x TBE buffer for 1 hour, results were visualized by Imaging System (Bio-Rad).

### Plaque assays

Plaque assays were performed as previously described^4, 30^. Briefly, reconstructed plasmid was transformed into *E. coli* DE3 competent cell. A single bacterial colony was picked from a fresh LB agar plate and grown in LB broth containing antibiotic at 37°C to an OD600 of ∼0.4. Protein expression was induced by the addition of 0.2 mM IPTG. After further growth for ∼3 h, 500 μl of the bacterial cultures was mixed with 14.5 ml of 0.5% LB top agar, and the entire samples were poured onto LB plates containing antibiotic and IPTG (0.1 mM). Plates were spotted with 4 μl of the T4 phage diluted in LB at eight 10-fold dilutions, namely, 10^0^-10^-7^. Lysate titer was determined using the small drop plaque assay method as previously described ^30, 31^. Plates were incubated at 37°C overnight and then imaged.

## Acknowledgments

Cryo-EM data of GajA was collected at OSU CEMAS with the assistance of Drs. Giovanna Grandinetti and Yoshie Narui. Cryo-EM data of GajAB was collected with the assistance of Drs. Adam D. Wier, Thomas J. Edwards, Tara Fox, and Jenny Wang at the National Cancer Institute Cryo-Electron Microscopy Center supported by grants from the NIH National Institute of General Medical Sciences (GM103310).

## Author contributions

T.M.F. conceived the project. X.Y.Y. and Z.S. performed molecular cloning, performed biochemical purification, nuclease activity, and plaque assays. Z.S. prepared EM grids, determined the cryo-EM structures. Q.L. helped on structural reconstruction. Z.S. and X.Y.Y. built the models. Z.S., X.Y.Y, and T.M.F. analyzed all the data together. T.M.F. wrote the manuscript with inputs from all the authors.

## Data and materials availability

Accession numbers for Gabija A tetramer (4A), Gabija AB complex 1 (4A:4B, C1 symmetry), and Gabija AB complex 2 (4A:4B, D2 symmetry) are as follows: (coordinates of atomic models: 8TK0, 8TK1, and 8TJY deposited to Protein Data Bank), and (density map: EMDB-41319, EMDB-41321, EMDB-41314, deposited to Electron Microscopy Data Bank.) All data needed to evaluate the conclusions are present in the paper.

---

**Extended data Fig. 1.**
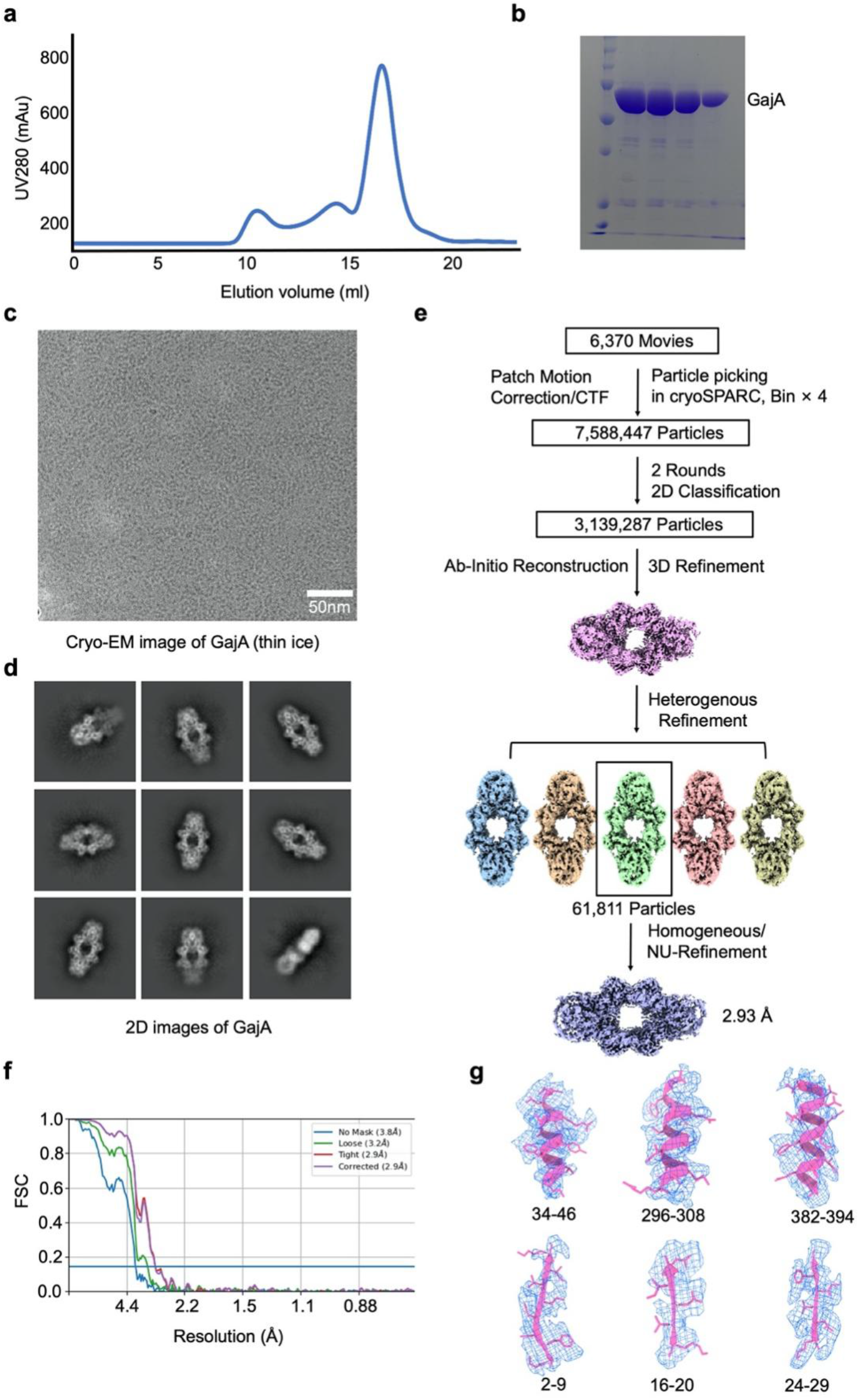
Cryo-EM reconstruction of GajA in thin ice. **a, b,** Gel filtration profile (**a**) and SDS-PAGE gel (**b**) of GajA purification. **c,** Cryo-EM image of GajA in thin ice. **d,** Representative 2D class averages of GajA calculated from thin-ice cryo-EM images. **e,** Data processing workflow for 3D reconstruction of GajA tetramer from thin-ice cryo-EM images. **f,** FSC curve of reconstructed GajA tetramer from thin-ice cryo-EM images. **g,** Representative cryo-EM density of GajA tetramer fit with α-helixes and β-strands. The density map is shown at contour levels of 0.03.

**Extended data Fig. 2.**
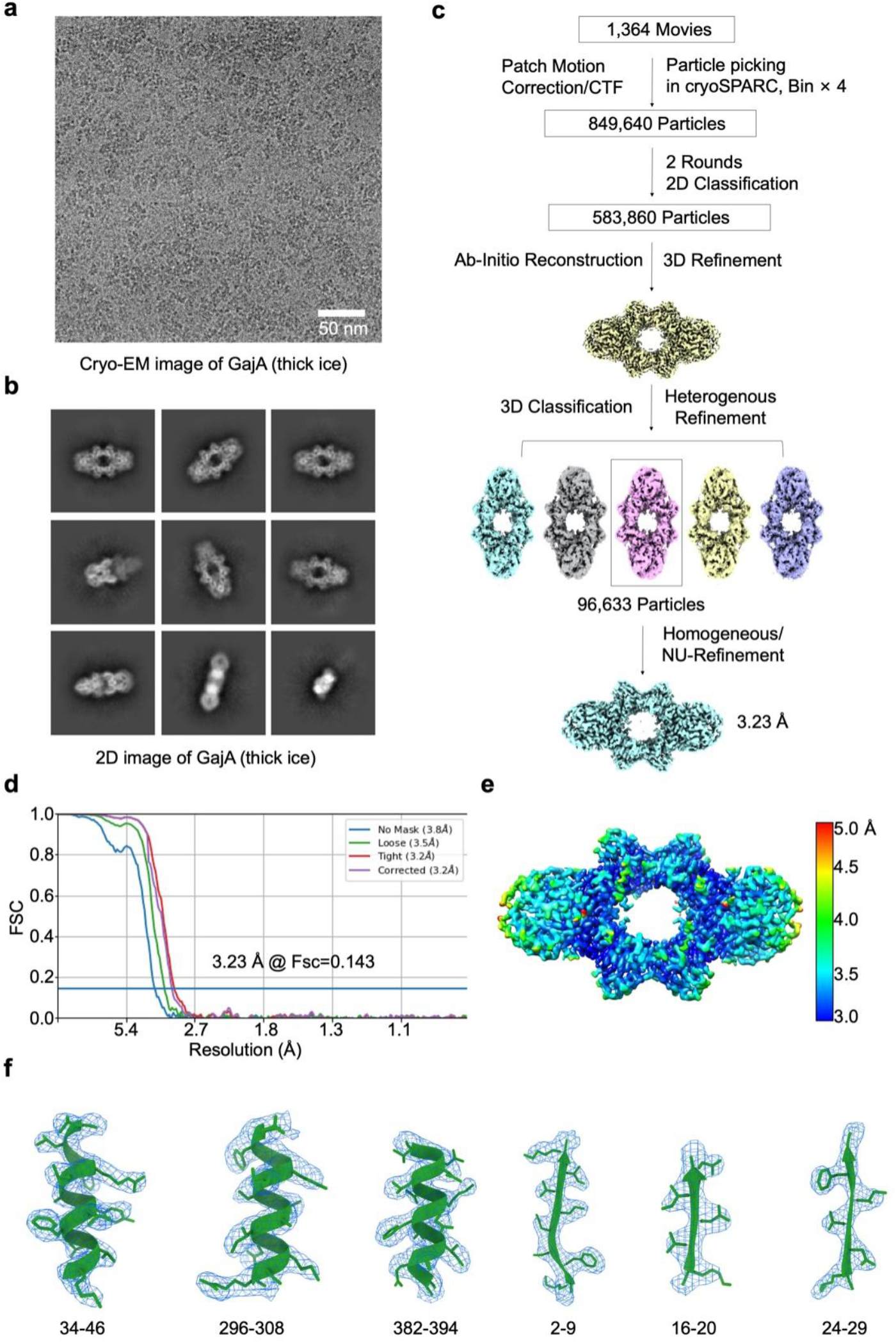
Cryo-EM reconstruction of GajA in thicker ice. **a,** Cryo-EM image of GajA in thick ice. **b,** 2D class averages of GajA calculated from thick-ice cryo-EM images. **c,** Data processing workflow for 3D reconstruction of GajA tetramer from thick-ice cryo-EM images. **d,** FSC curve of reconstructed GajA tetramer from thick-ice cryo-images. **e,** Local resolution of reconstructed GajA tetramer from thick-ice cryo-images. **f,** Cryo-EM density of GajA tetramer fit with α-helixes and β-strands. The density map is shown at contour levels of 0.03.

**Extended data Fig. 3.**
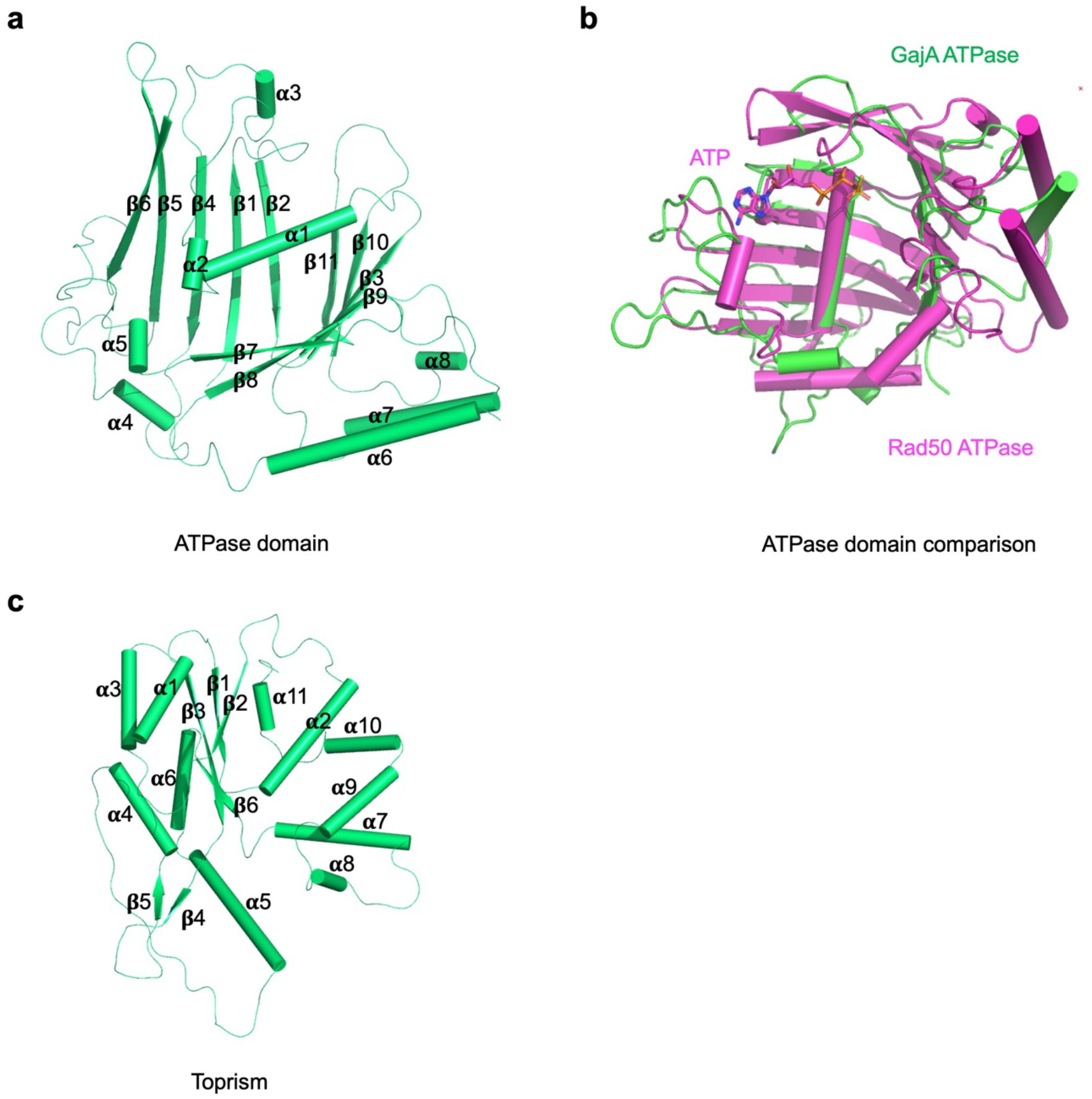
Architecture of GajA. **a,** Ribbon diagram of GajA N-terminal ATPase domain with secondary structures indicated. **b,** Overlaid structures of GajA N-terminal ATPase domain (green) and Rad50 ATPase domain (PDB ID 5DNY, magenta). **c,** Ribbon diagram of GajA C-terminal Toprim domain with secondary structures indicated.

**Extended data Fig. 4.**
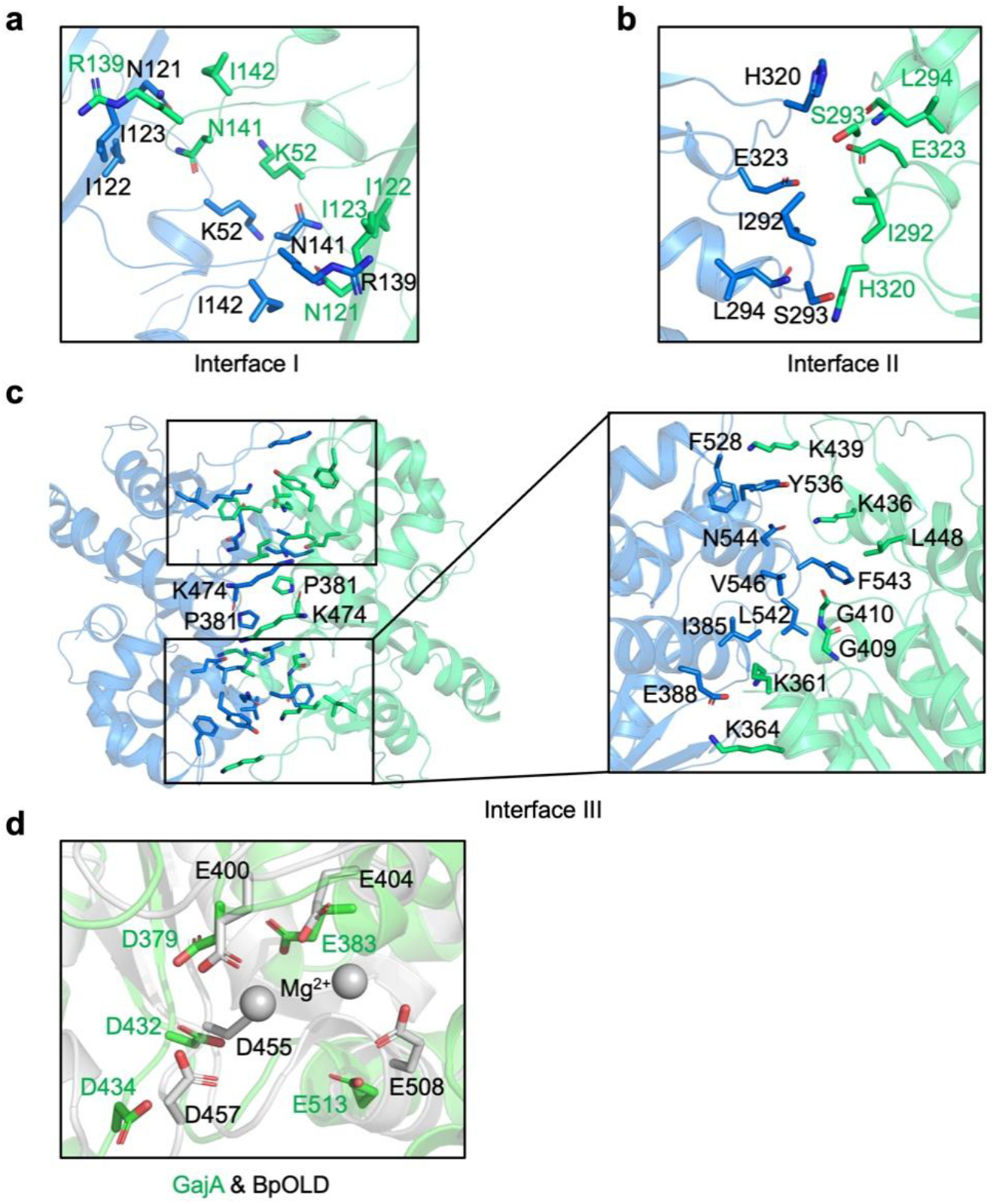
Interfaces in GajA tetramer. **a-c,** Enlarged views of interface I (**a**), interface II (**b**), and interface III (**c**) in GajA tetramer. Key residues on the interfaces were highlighted in sticks. **d,** Superimposed structures of the active sites from GajA (green) and BpOLD (PDB ID 6NK8, grey).

**Extended data Fig. 5.**
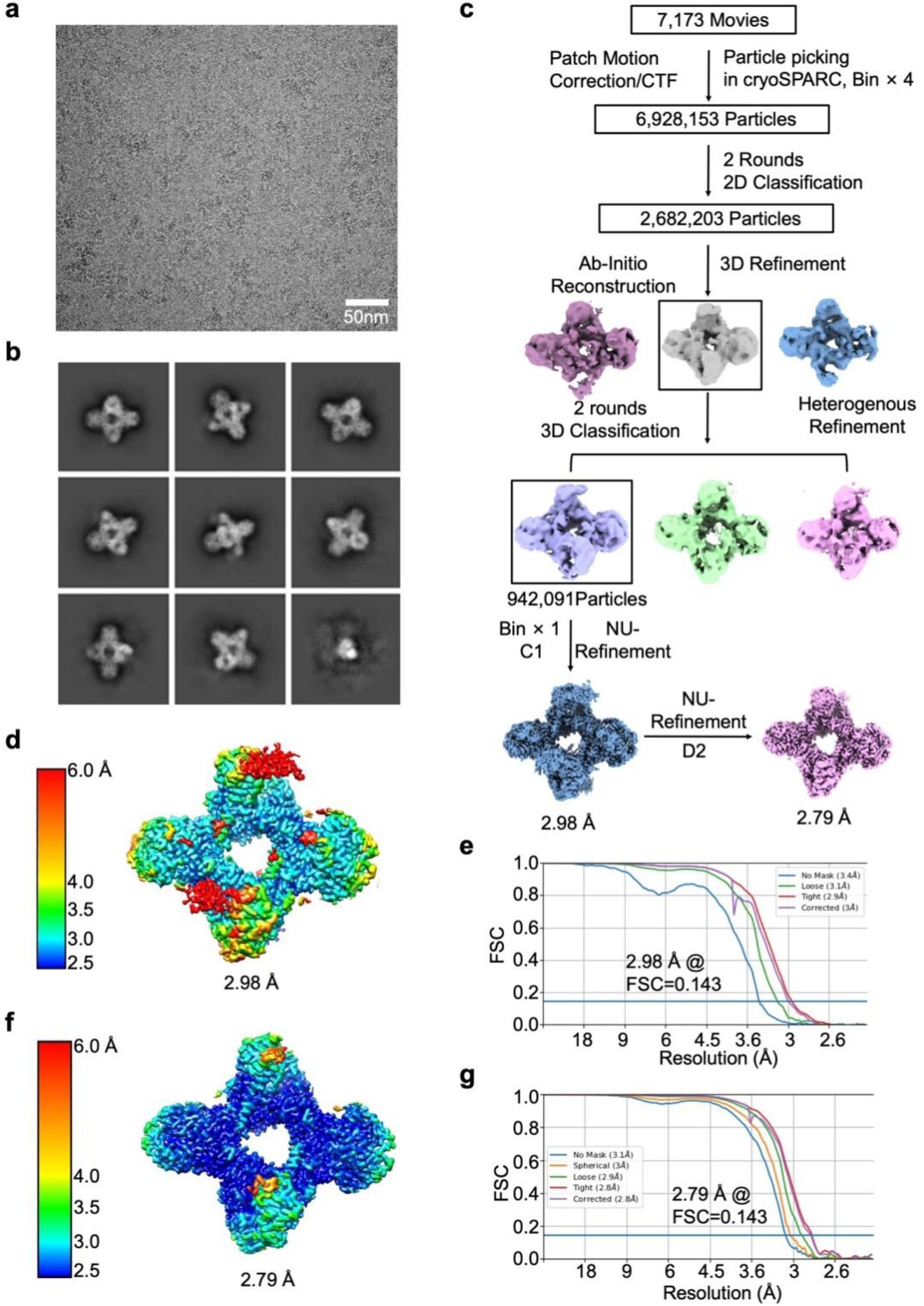
Cryo-EM reconstruction of GajAB. **a,** Cryo-EM image of GajAB complex. **b,** 2D class averages of GajAB complex. **c,** Data processing workflow for 3D reconstruction of GajAB complex. **d, e,** Local resolution (**d**) and FSC curve (**e**) of reconstructed GajAB complex without symmetry setting. **f, g,** Local resolution (**f**) and FSC curve (**g**) of reconstructed GajAB complex with D2 symmetry setting.

**Extended data Fig. 6.**
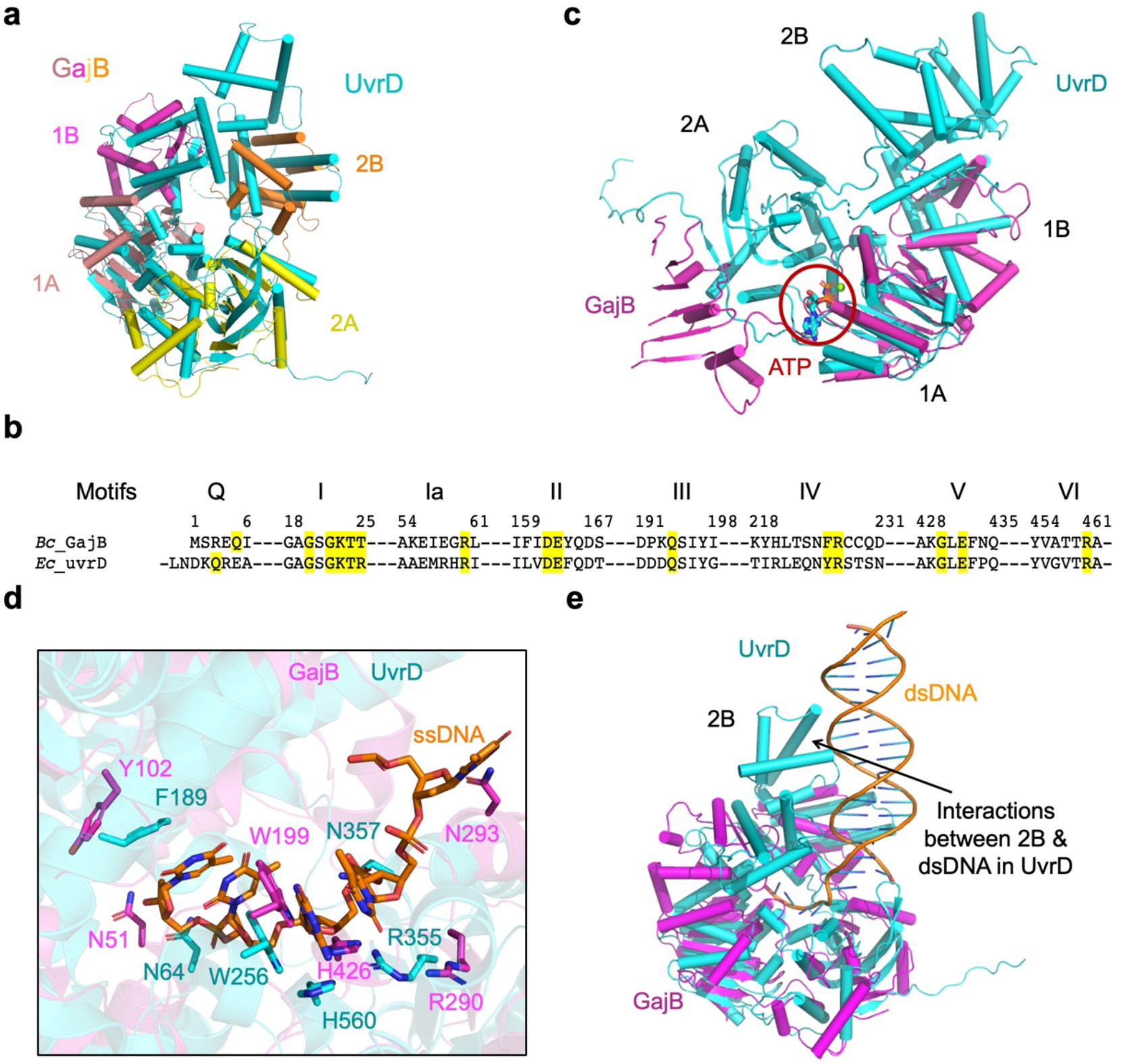
Structural comparison of GajB and UvrD. **a,** Overlaid structures of GajB (magenta, pink, yellow, and orange) and UvrD (PDB ID 2IS2, blue). **b,** Sequence alignment of ATP binding motifs between GajB and UvrD. **c,** Overlaid structures of GajB (magenta) and UvrD (blue) showed that domain 2A of GajB is not well positioned to coordinate ATP. **d,** Expanded view of key residues involved in coordinating ssDNA from GajB (magenta) and UvrD (blue). **e,** Overlaid structures of GajB (magenta) and UvrD (blue) revealed that domain 2B in GajB lacks key motifs for coordinating dsDNA.

**Extended data Fig. 7.**
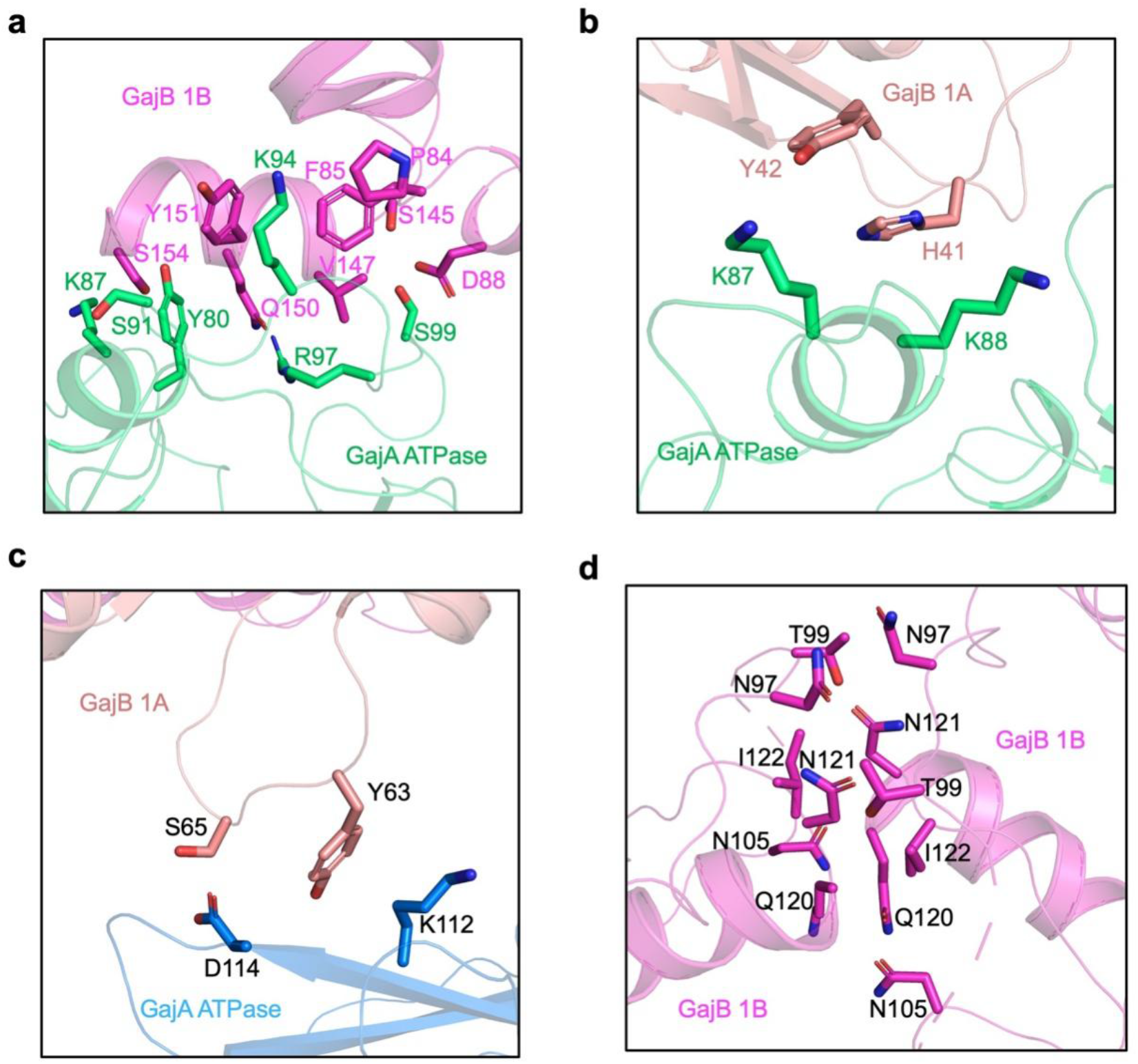
Interfaces in GajAB. **a,** Key residues mediating interactions between GajB domain 1B (magenta) and GajA ATPase domain (green). **b,** Key residues mediating *cis*-interactions between GajB domain 1A (pink) and GajA ATPase domain (green). **c,** Key residues mediating trans-interactions between GajB 1A (pink) and GajA ATPase (blue). **d,** Key residues mediating interactions of two neighboring GajB protomers.

**Extended Data Table 1.** Cryo-EM data collection, refinement, and validation statistics.

|  | Gabija A tetramer<br>(4A)<br>(EMDB-41319)<br>(PDB 8TK0) | Gabija AB complex 1<br>(4A:4B, C1 symmetry)<br>(EMDB-41321)<br>(PDB 8TK1) | Gabija AB complex 2<br>(4A:4B, D2 symmetry)<br>(EMDB-41314)<br>(PDB 8TJY) |
| --- | --- | --- | --- |
| **Data collection and processing** |  |  |  |
| Magnification | 81,000 | 81,000 | 81,000 |
| Voltage (kV) | 300 | 300 | 300 |
| Electron exposure (e-/Å<sup>2</sup>) | 50 | 50 | 50 |
| Defocus range (μm) | 1.0 - 2.0 | 0.5 - 2.0 | 0.5 - 2.0 |
| Pixel size (Å) | 0.899 | 1.12 | 1.12 |
| Symmetry imposed | D2 | C1 | D2 |
| Initial images (no.) | 1,364 | 7,173 | 7,173 |
| Initial particle images (no.) | 849,640 | 6,928,153 | 6,928,153 |
| Final particle images (no.) | 96,633 | 942,091 | 942,091 |
| Map resolution (Å) | 3.23 | 2.98 | 2.79 |
| FSC threshold | 0.143 | 0.143 | 0.143 |
| Map resolution range (Å) | 3.23 - 5.0 | 2.98 - 6.0 | 2.79 - 6.0 |
| **Refinement** |  |  |  |
| Initial model used (PDB code) | AlphaFold | AlphaFold | AlphaFold |
| Model resolution (Å) | 3.2 | 3.1 | 3.1 |
| FSC threshold | 0.5 | 0.5 | 0.5 |
| Map sharpening *B* factor (Å<sup>2</sup>) | -141.5 | -110.9 | -121.5 |
| Model composition |  |  |  |
| Non-hydrogen atoms | 14816 | 23424 | 22692 |
| Protein residues | 1816 | 2877 | 2788 |
| Ligands | 0 | 0 | 0 |
| Nucleotide | 0 | 0 | 0 |
| *B* factors (Å<sup>2</sup>) |  |  |  |
| Protein | 78.80 | 47.07 | 45.75 |
| Ligand |  |  |  |
| Nucleotide |  |  |  |
| R.m.s. deviations |  |  |  |
| Bond lengths (Å) | 0.004 | 0.004 | 0.003 |
| Bond angles (°) | 0.614 | 0.938 | 0.469 |
| Validation |  |  |  |
| MolProbity score | 1.71 | 1.40 | 1.40 |
| Clashscore | 5.33 | 3.42 | 3.34 |
| Poor rotamers (%) | 0.00 | 0.08 | 0.00 |
| Ramachandran plot |  |  |  |
| Favored (%) | 93.61 | 96.12 | 96.01 |
| Allowed (%) | 6.39 | 3.88 | 3.99 |
| Disallowed (%) | 0.00 | 0.00 | 0.00 |

